# Infernal-GPU: accurate E-values and sensitive covariance-model search for structured RNAs

**DOI:** 10.64898/2026.11.15.766754

**Authors:** Ding Wang, Junru Jin, Jianbo Qiao, Leyi Wei, Shu Wu, Qiang Liu

## Abstract

**Motivation:** Covariance models (CMs) find structured RNAs by scoring sequence and base pairing together, and they underlie the Infernal software and the Rfam database. Their dynamic programming is expensive, so Infernal relies on two approximations: E-values are calibrated on a short stretch of random sequence, and fast sequence-only filters decide which parts of a target the CM examines. How much accuracy these approximations cost, and where, is largely unknown, because removing them on CPUs is prohibitively slow.

**Results:** We present Infernal-GPU, a GPU implementation of Infernal’s CM dynamic programming that matches Infernal’s results. Its key step evaluates the one serial recurrence of CM scoring, the self-loop of insert-left states, as two parallel prefix scans. One GPU calibrates all 4,227 Rfam models overnight, which makes calibrations on ten times more random sequence routine. These longer calibrations show that the E-values distributed with Rfam are off by more than twofold for one model in eight. An exhaustive search reference shows that sequence filters discard remote homologs whose similarity lies almost entirely in base pairing; behind permissive filters, Infernal-GPU recovers them with a third of the CPU work that Infernal needs at the same setting.

**Availability and implementation:** Infernal-GPU (CUDA, C++ and Python) is freely available at https://github.com/void-echo/infernal-gpu and from PyPI (`pip install infernal-gpu`). Rfam 15.0 models recalibrated on ten times more random sequence are attached to the software release.

**Supplementary information:** Supplementary data are available online.

## 1 Introduction

Many functional RNAs conserve their secondary structure long after their sequence has diverged, so their distant relatives are often invisible to sequence comparison (Freyhult *et al*. 2007). Covariance models, stochastic grammars that score a sequence and its base pairs together, were developed to find such relatives (Eddy and Durbin 1994, Sakakibara *et al*. 1994, Durbin *et al*. 1998). Through Infernal (Nawrocki *et al*. 2009, Nawrocki and Eddy 2013), CMs define the families of the Rfam database (Ontiveros-Palacios *et al*. 2025) and annotate non-coding RNAs in genomes and metagenomes (Kalvari *et al*. 2021). New models are built continually: at each Rfam release, in RNA motif discovery (Yao *et al*. 2006, Weinberg *et al*. 2017), and for every query in the alignment pipelines that feed RNA structure predictors such as trRosettaRNA and RhoFold+ (Zhang *et al*. 2021, 2023, Wang *et al*. 2023, Shen *et al*. 2024).

CM dynamic programming grows with the number of model states, the target length and the range of subsequence lengths, even when query-dependent bands restrict the lengths each state considers (Nawrocki and Eddy 2007). Infernal makes it practical through two approximations. In search, profile hidden Markov model (HMM) filters that ignore base pairing discard most of the target before the CM sees it, which made Infernal 1.1 about a hundred times faster than its predecessor (Weinberg and Ruzzo 2006, Eddy 2011, Kolbe and Eddy 2011, Nawrocki and Eddy 2013). For statistics, cmcalibrate scores 1.6 Mb of random sequence with the complete model in four modes and fits exponential tails to the random hits (Klein and Eddy 2003, Nawrocki and Eddy 2013); no filter can be used, because the quantity measured is the model’s own behaviour on random sequence. Even this short calibration takes hours per model on one CPU (Nawrocki and Eddy 2023), about 2,800 CPU core-hours for the Rfam library, and it lies on the critical path of pipelines such as rMSA and RNAcmap that calibrate a new model for each query (Zhang *et al*. 2021, 2023). GraphClust2 shortens it further, accepting less reliable E-values (Miladi *et al*. 2019).

The cost of these approximations is only partly known, because removing them is expensive. The overall loss of sensitivity to the filters was measured when they were introduced (Nawrocki and Eddy 2013), but which homologs they discard, and why, has not been examined, and the sampling error of calibration has not been measured at all. Both questions require the full CM computation at a scale that CPUs make impractical: exhaustive searches, and calibrations many times longer than the default.

Graphics processing units (GPUs) have transformed protein homology search (Khajeh-Saeed *et al*. 2010, Jiang and Ganesan 2016, Schmidt *et al*. 2024, Kallenborn *et al*. 2025). For RNA, a GPU sequence prefilter has been combined with CM realignment on the CPU (Park *et al*. 2026), and GPUs accelerate secondary-structure prediction (Rizk and Lavenier 2009, Lei *et al*. 2012, Stern and Mathews 2013) and the parsing of stochastic grammars (Johnson 2011, Yi *et al*. 2014), where all cells of one span are independent. Scanning genomes with CMs poses two further obstacles: insert-left states loop on themselves, so each subsequence length waits for the previous one, and per-state bands make the working memory large and irregular. Hardware implementations on field-programmable gate arrays have accelerated CM alignment and Infernal’s filters (Moscola *et al*. 2010, Xia *et al*. 2010, McVicar *et al*. 2013), but not calibration.

Here we present Infernal-GPU, a GPU implementation of Infernal’s CM dynamic programming. It parallelizes the insert-left recurrence exactly, stores only the cells inside each state’s band and lets Infernal compute the final scores and E-values of every search. We use it to measure both approximations and to remove them: it calibrates the Rfam library overnight, quantifies the sampling error of current E-values (Fig. 1A) and makes ten-fold longer calibrations practical, and it provides an exhaustive search reference that identifies the homologs the filters lose, shows why they are lost and recovers them behind permissive filters.

**Fig. 1.**
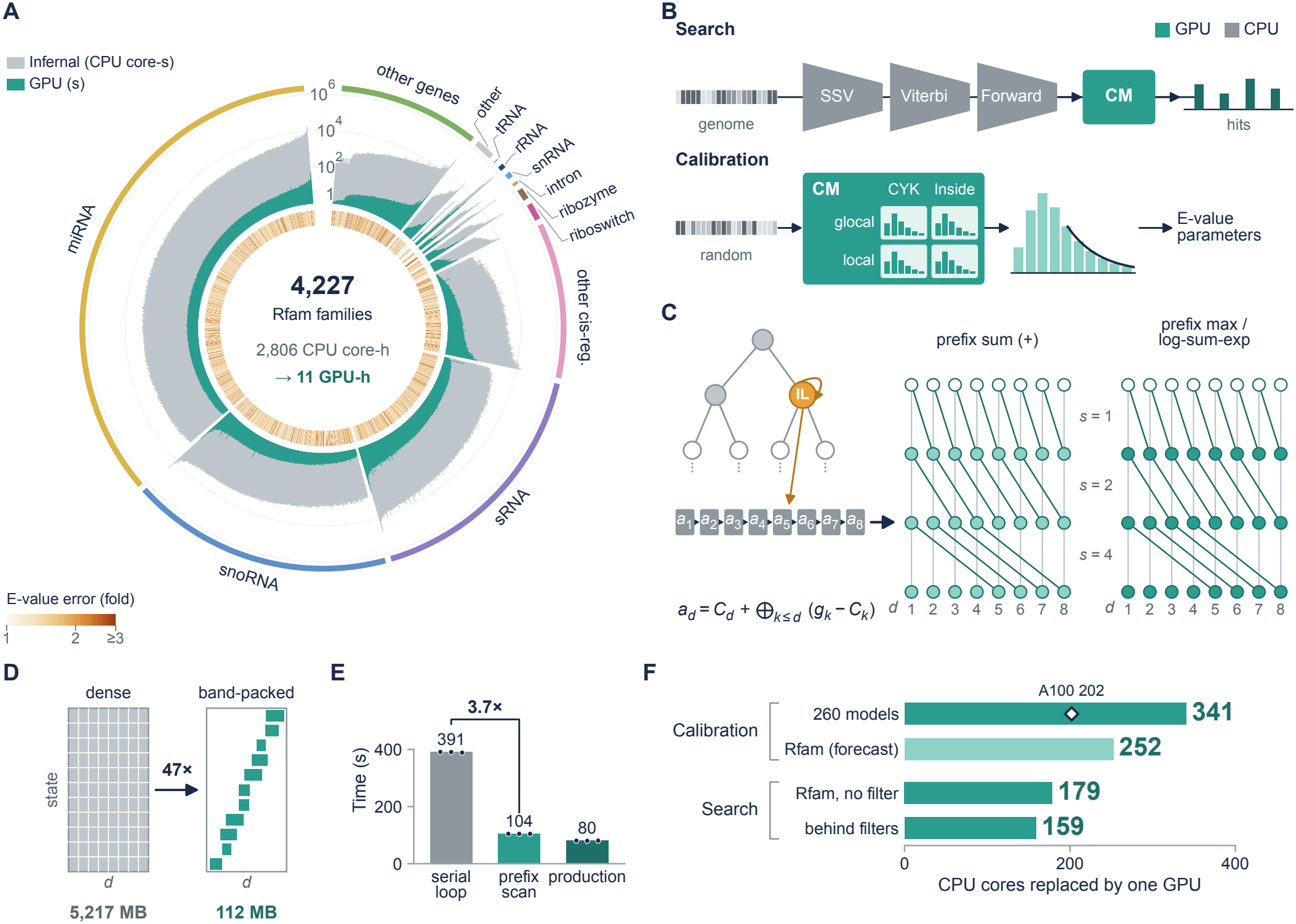
Infernal-GPU computes covariance-model dynamic programming on a GPU. (**A**) The Rfam 15.0 library: its 4,227 models grouped by RNA class (outer arcs), each with its cmcalibrate CPU core-seconds (forecast, grey) and GPU seconds (teal) on a logarithmic radial scale, and the E-value error of the calibration distributed with Rfam (local Inside, against a 16-Mb GPU calibration; inner ring). (**B**) Infernal runs full CM dynamic programming behind its profile-HMM filters during search and on random sequence in four modes during calibration; Infernal-GPU computes these stages on the GPU (teal). (**C**) An insert-left state (amber) loops on itself, so its score at each length waits for the previous one; the chain equals an emission prefix sum followed by a prefix maximum (CYK) or log-sum-exp (Inside), evaluated for all lengths in logarithmic depth. (**D**) Storing only each state’s band reduces the per-window workspace of the largest Rfam model 47-fold. (**E**) End-to-end time of an eight-model search of the *M. ruminantium* genome with serial and prefix evaluation of insert-left states (same storage and tiling) and in the production configuration; bars, medians of three runs (points). (**F**) CPU cores of Infernal work taken over by one GPU (bars, RTX 4090) in calibration (260 benchmark models, measured; Rfam library, forecast) and search (filter-free search of the Rfam library; CM stages behind Infernal’s most permissive filters); diamond, the same calibration on an A100 GPU.

## 2 Methods

### 2.1 Covariance-model dynamic programming

A CM is a tree of states of nine types (Eddy and Durbin 1994, Durbin *et al*. 1998). For state v, end position j and subsequence length d, the CYK score is the maximum, and the Inside score the log_2_-sum-exp, over v’s children of child score plus transition score, plus the emission score of the residues that v emits; bifurcations combine their two children over all splits of d. Query-dependent banding restricts each state to a length interval (Nawrocki and Eddy 2007); Infernal uses tight bands (β_1_) for its CYK filter and looser bands (β_2_) for Inside and calibration, and the maximum hit length W bounds all bands. We follow Infernal 1.1.5 in all scoring details, including local begins and ends and NULL3 composition correction. Infernal does not search for truncated hits in its filter-free mode or whenever it scores with QDB bands, so all searches, by Infernal and by Infernal-GPU, used its non-truncated mode (`--notrunc`) to keep settings comparable.

### 2.2 Prefix-scan evaluation of insert-left states

A state’s score at (j, d) combines the scores of its children, which are computed before it, except where an insert state loops on itself. An insert-right state emits residue j, so its self-loop reads its score at the previous end position j − 1, which is already complete; an insert-left state emits the leftmost residue j − d + 1, so its self-loop reads its own score at the same j and length d − 1. This is the only dependency within a state and end position.

At a fixed insert-left state and end position, let a_d_ be its score at length d, e_d_ its emission score, t its self-transition score and n_d_ the ⊕-combination of all other child contributions, where ⊕ is maximum (CYK) or log_2_-sum-exp (Inside). The recurrence a_d_ = (n_d_ ⊕ (a_d−1_ + t)) + e_d_ makes each length wait for the previous one. Let g_d_ = n_d_ + e_d_, with any lower-boundary term folded into the first length d_0_ of the band, and C_d_ = Σ_r=d0..d_ e_r_ + (d − d_0_)t. Entering the loop at length k and staying to length d adds exactly C_d_ − C_k_ to g_k_, so a_d_ = C_d_ + ⊕_k=d0..d_ (g_k_ − C_k_).

The recurrence thus reduces to one prefix sum followed by one prefix maximum or prefix log_2_-sum-exp, associative scans that p threads evaluate in O(log p) steps (Kogge and Stone 1973, Ladner and Fischer 1980, Blelloch 1990). Threads of a CUDA block are assigned to lengths; warp-shuffle scans combine within warps, shared memory across warps, and longer bands are processed in blocks with carried totals. The closed form is exact in real arithmetic; in floating point it changes only the order of operations, and parallel and serial evaluation differed by at most 1.2 × 10^−4^ bits. Differences from Infernal’s own scores near the reporting thresholds are larger (Supplementary Fig. S1) because Infernal evaluates log-sums with a lookup table.

### 2.3 GPU implementation

Each CUDA block processes one sequence window, with its threads assigned to sub-sequence lengths, and traverses states and end positions in dependency order; a C++ template selects the combination operator, so CYK and Inside share one code path. For a bifurcation, each thread combines the scores of all splits of its length (maximum for CYK, log-sum for Inside), reading the right child at the current end position and the left child at earlier end positions from a ring buffer. Each state stores only the cells of its band, and the ring buffers keep only the lengths that the parent bifurcation reads. Windows overlap by W residues; their length and the storage layout are chosen per model from device memory, and the number of windows per launch is set so that the last wave of blocks fills the GPU. NULL3 corrections and score gates are computed in the root-state pass. The production configuration combines the prefix scan with this wave-aware tiling and band-packed storage.

### 2.4 Calibration

Infernal-GPU follows cmcalibrate. The random sequence is 160 chunks of 10 kb sampled from Infernal’s built-in genomic HMM with seed 181, reset before each model, generated with Infernal’s own routines (other seeds and the 16-Mb sequences likewise); the four modes (glocal and local alignment, each scored with CYK and Inside) use β_2_ bands; and exponential tails are fitted as in cmcalibrate and written into the model file. LSU_rRNA_eukarya was calibrated with tailn = 200, as its distributed parameters indicate.

For each end position j, Infernal’s greedy rule visits lengths in increasing order and reports a hit when its score, before and after NULL3 correction, exceeds the corrected score of the last hit reported at j; reported hits are then filtered for overlap. Because the NULL3 correction is always positive, a cell is reported exactly when its corrected score passes the cutoff and exceeds the corrected scores of all shorter lengths at j. With the same prefix machinery, the GPU computes this running maximum m(d) over all shorter lengths and emits cell (j, d) only if its corrected score exceeds m(d) − δ, with δ = 0.05 bits. Host and GPU corrected scores differ by about 10^−3^ bits, far below δ/2, so every cell that the rule could report is emitted, and applying the rule on the host yields exactly the random hits obtained from all cells.

Calibrations are compared by x_0.01_, the bit score at which E = n_fit_ (Z/L) exp(−λ(x − μ)) equals 0.01 in a search of Z = 100 Mb, with each calibration’s own number of fitted hits n_fit_ and random-sequence length L. For the glocal modes of five giant models (λ < 0.05), x_0.01_ is dominated by the printed precision of λ; there we compared λ (agreement within 3 × 10^−5^) and random-hit counts. Sampling error was measured by recalibrating the 260 benchmark models (Section 2.6) with other seeds and on 16 Mb (seeds 181 and 1), and accuracy against the independent 16-Mb calibration (seed 1). The E-value error of the distributed Rfam parameters is the fold difference between 0.01 and the E-value they assign to the score at which the 16-Mb calibration of the library (seed 181) gives E = 0.01. Whole-library CPU cost is cmcalibrate’s `--forecast` estimate, whose median ratio to measured cost on the benchmark models was 1.07.

### 2.5 Search

In filter-free search, the GPU runs CYK with β_1_ bands, gated at the calibrated threshold of Infernal’s CYK filter (P = 10^−4^) on NULL3-corrected scores, and then Inside with β_2_ bands on the surviving intervals, gated at the requested E-value. Both gates are lowered by 0.25 bits, well beyond the largest GPU underestimate of Infernal’s scores near the thresholds (Supplementary Fig. S1). cmsearch then rescans the candidate intervals, padded by W, with the original search space and reports the final hits; a streaming driver overlaps these rescans with the GPU scan of the next model. Filter-free search reproduces `cmsearch --nohmm`. Exhaustive search, which also skips the CYK gate, reproduces `--nohmm --noF6`: it applies no filter and scores every position with the QDB-banded CM, whose bands, as in Infernal’s filter-free mode, are bounded by W.

Behind the HMM filters, a small opt-in hook in Infernal’s search pipeline (added to a separate Infernal 1.1.5 build by `infernal-gpu patch-infernal`) writes out the envelopes that pass the filters instead of scoring them in the CM stages; the GPU scores the envelopes with QDB Inside, and cmsearch rescans the candidates. Filter settings were default, `--mid` and `--mid --Fmid 0.1`. The factorial design crossed the early filters (kept, or SSV and Viterbi with its bias filter skipped) with Forward-stage thresholds of 0.003, 0.02 and 0.1, applied together to F3–F5 and their bias filters. Replays kept the exhaustive hits inside the envelopes passed by each setting.

### 2.6 Benchmarks and evaluation

The remote-homology benchmark follows Infernal’s RMARK procedure (Nawrocki and Eddy 2013) on Rfam 15.0 seed alignments: test sequences with less than 60% identity to the training set are embedded in about 20.8 Mb of synthetic background, and models are built and calibrated from the training alignments. Leaving out SSU_rRNA_eukarya, whose model alone would cost several times the rest of the benchmark in Infernal’s filter-free runs, gives 260 families and 1,756 positives. Three replicates vary the test representatives, background and embedding; two further benchmarks used background derived from real sequence (dinucleotide-preserving shuffles of 1.2-kb fragments of the *Escherichia coli* genome and a human gut metagenome assembly) and a 55% identity split (150 families, 881 positives). Hits were classified as in the RMARK scripts, and we report sensitivity at one false positive per query with 95% intervals from 1,000 bootstrap resamples of families, paired across methods (Efron and Tibshirani 1993). A positive counts as found when a true-positive hit has E ≤ 0.01.

Each positive was also cut out in its own orientation and scored without filters by its family’s profile HMM alone (`cmsearch --hmmonly`, all thresholds 1, no bias filter), by its CM (`--nohmm --noF6`) and by a structure-free CM built from the same alignment with `cmbuild --noss` and calibrated with Infernal-GPU. The base-pair contribution is the CM minus the structure-free CM score, and its share is this difference divided by the CM score (positives with CM scores of at least 10 bits). RNA classes follow the Rfam type annotation. The genome search used *Methanobrevibacter ruminantium* M1 (NC_013790.1) (Leahy *et al*. 2010) with all 4,227 Rfam 15.0 models.

### 2.7 Timing

GPU times are elapsed times on one NVIDIA RTX 4090 in a dual-socket AMD EPYC 7742 server, on otherwise idle GPUs; CPU work is user plus system time. CPU-core equivalents are the Infernal CPU work that the GPU takes over, minus that of the remaining rescans, divided by GPU time. Fixed-allocation runs used 32 cores (`cmcalibrate --cpu 32`, or 32 single-threaded search processes plus one GPU); node time, CPU work divided by 32 plus GPU time, matched them within 8%.

## 3 Results

### 3.1 A prefix formulation puts CM dynamic programming on the GPU

Infernal must run the full CM dynamic programming in two places: behind its filters during search and on random sequence during calibration (Fig. 1B). For every state and end position, this computation scores all subsequence lengths, which are independent at every state type except insert-left, whose self-loop makes the score at length d depend on that at d − 1 (Fig. 1C). Unrolled, the chain is an emission prefix sum followed by a prefix maximum for CYK or a prefix log-sum-exp for Inside, which a block of threads computes in logarithmic depth (Section 2.2). Parallel scans have broken sequential dependencies in GPU sequence alignment and HMM inference (Ganesan *et al*. 2010, Khajeh-Saeed *et al*. 2010, Hassan *et al*. 2021); here the dependency lies inside a stochastic grammar and is resolved for maximization and summation alike, so CYK and Inside share one implementation.

With storage and tiling held fixed, the prefix scan alone sped up a search of eight Rfam models against the 2.94-Mb *M. ruminantium* genome 3.7-fold, and the production configuration, which adds wave-aware tiling and band-packed storage, a further 1.3-fold (Fig. 1E). Band-packed storage reduced the per-window memory of the largest Rfam model, the eukaryotic large-subunit rRNA, from 5.2 GB to 112 MB (Fig. 1D). Across calibration and search, one RTX 4090 took over the CM work of 160–340 CPU cores (Fig. 1F). The final hits come from Infernal’s rescans of the GPU candidates: a filter-free search of all 4,227 Rfam models against the *M. ruminantium* genome returned the same 129 hits as `cmsearch --nohmm`, in 4.8 h on one GPU against 851 CPU core-hours, and searches of the 260 benchmark families returned exactly Infernal’s hits (Supplementary Table S1).

### 3.2 One GPU calibrates the Rfam library overnight

Infernal-GPU reproduces the cmcalibrate protocol step by step, with the same random sequence, modes, bands, hit-collection rule and fits (Section 2.4), and its calibrations agreed closely with Infernal’s. For the 4,227 Rfam models, the fitted exponential slopes λ and the numbers of random hits differed from those distributed with Rfam by a median of 0.08% and 0.02% (Fig. 2B, C). On identical random sequence, the E = 0.01 score threshold differed between GPU and Infernal calibrations by a median of 0.023 bits, whereas recalibration with another random seed moved it by 1.0 bit, 44 times more (Fig. 2D); in every mode and range of model size, seed-to-seed noise exceeded the GPU–Infernal difference 20- to 88-fold (Fig. 2E). The parameters were bit-identical on a consumer and a data-centre GPU (Supplementary Table S1), and models calibrated on the GPU returned the same benchmark hits as models calibrated by Infernal.

**Fig. 2.**
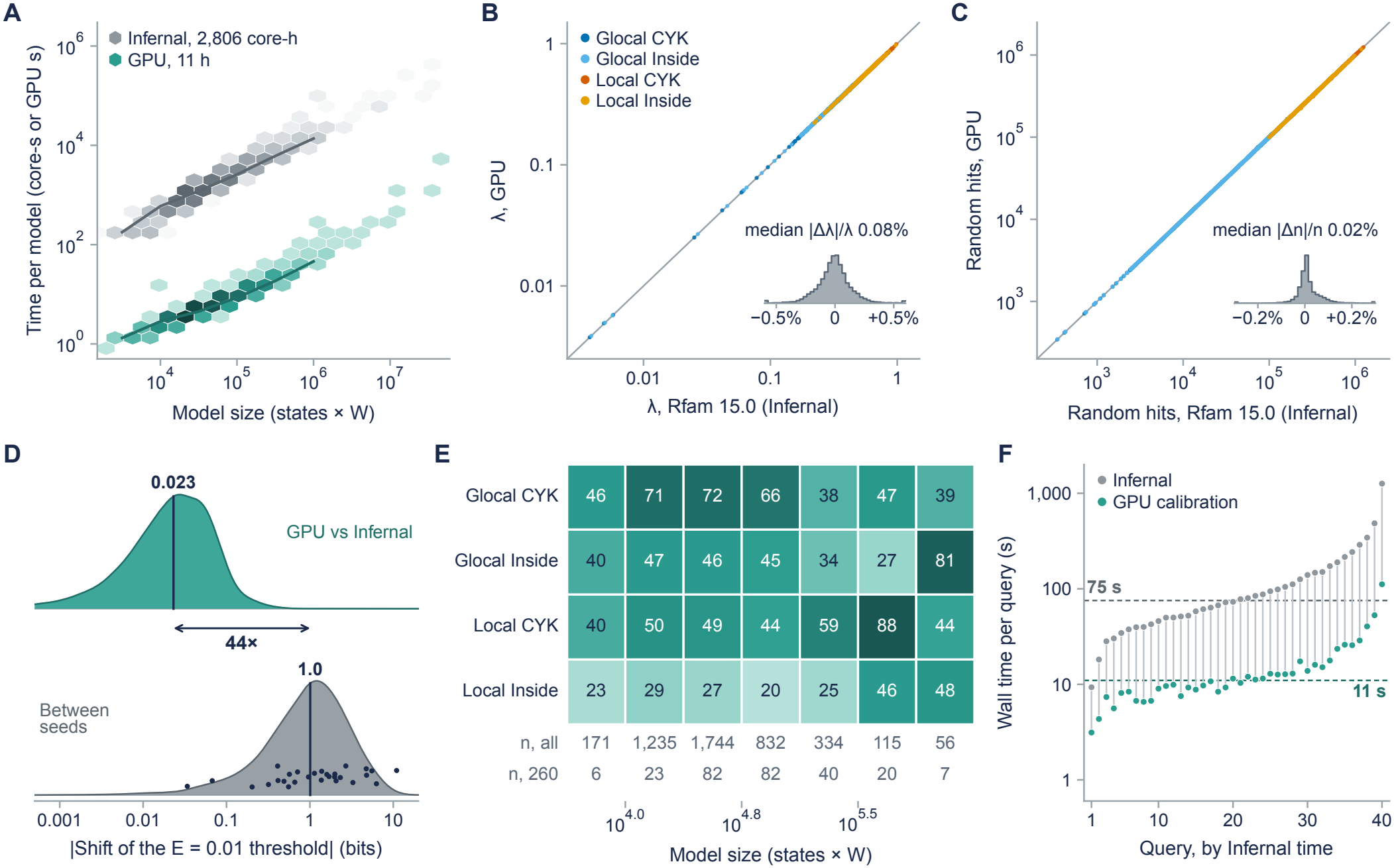
One GPU calibrates the Rfam library overnight with Infernal’s results. (**A**) Calibration time per model against model size (states × maximum hit length W) for the 4,227 Rfam models: cmcalibrate CPU core-seconds (forecast) and GPU seconds; lines, binned medians. (**B, C**) Exponential slope λ (**B**) and number of random hits (**C**) of GPU calibrations against those distributed with Rfam 15.0, for 4,227 models in four modes; insets, relative differences. (**D**) Shift of the bit score giving E = 0.01 in a 100-Mb search between GPU and Infernal calibrations on identical random sequence (260 benchmark and 4,227 Rfam models, four modes) and between random seeds (260 models; points, seven models recalibrated by cmcalibrate). (**E**) Ratio of the seed-to-seed shift to the GPU–Infernal shift (medians) by mode and model size; numbers of models per size bin below. (**F**) Wall time per query for building, calibrating and searching with a new model on one 32-core server, with cmcalibrate and with GPU calibration, for 40 benchmark families spanning the range of calibration cost; dashed lines, medians.

Calibration fell from CPU-hours to seconds or minutes per model, across every class of RNA in Rfam (Figs 1A and 2A). The Rfam library took 11 h on one GPU against about 2,800 CPU core-hours, and its largest model, quoted at about 30 h on 12 threads (Nawrocki and Eddy 2023), took 73 min. Against cmcalibrate on a 32-core server, one GPU was 8–13 times faster for models ranging from a selenocysteine insertion element to bacterial small-subunit rRNA (Supplementary Table S2). The gain matters most where a model is built for every query: in a workflow that builds a CM, calibrates it and searches a 20.8-Mb database on a 32-core server, calibration took 99% of Infernal’s time, and GPU calibration shortened the median query from 75 to 11 s (Fig. 2F).

### 3.3 Default calibrations leave E-values with a large random error

The tail that a calibration fits depends on which 1.6 Mb of random sequence was drawn, so calibrations of one model with different seeds fan out around the significance threshold (Fig. 3A), and the difference reaches users as noise in the E-values of their hits. When the benchmark models were recalibrated with another seed, the E-values of the same hits changed by a median of 1.3-fold, and by more than 2.2-fold in one family in twenty (Fig. 3D), and up to ten true homologs crossed cmsearch’s inclusion threshold of E = 0.01. Between GPU and Infernal calibrations on the same random sequence, none did.

**Fig. 3.**
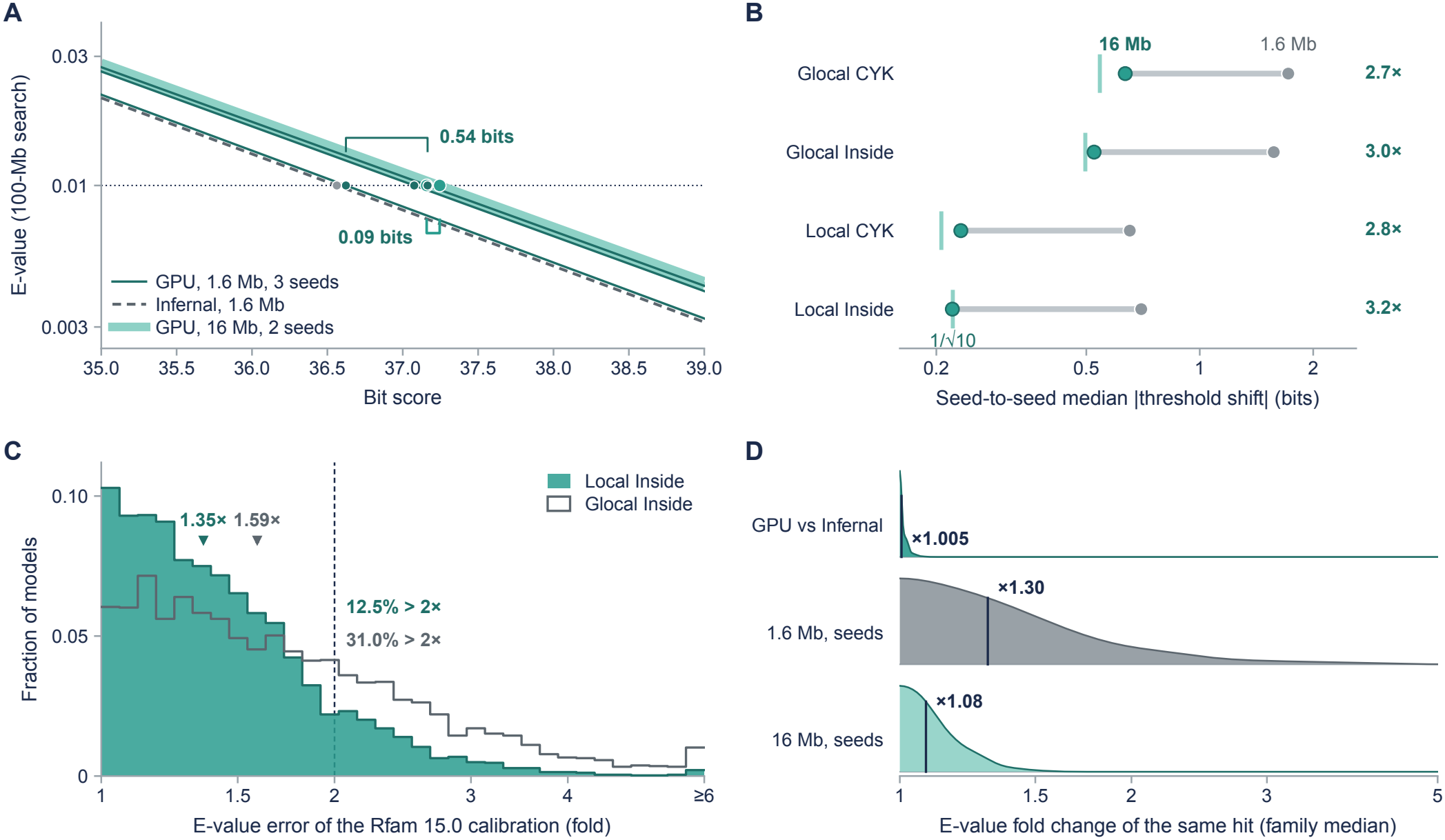
Default calibrations leave E-values with a large random error, which ten-fold longer calibrations reduce about threefold. (**A**) E-value in a 100-Mb search against bit score for a benchmark model of median calibration cost (DUF3800-I, local Inside), calibrated on the default 1.6 Mb by Infernal (seed 181) and by the GPU (seeds 181, 1 and 2), and on 16 Mb by the GPU (seeds 181 and 1). (**B**) Seed-to-seed shift of the E = 0.01 threshold by mode at 1.6 and 16 Mb (260 models); ticks, the 1/√10 expectation. (**C**) E-value error of the calibrations distributed with Rfam 15.0 against 16-Mb GPU calibrations of all 4,227 models, local and glocal Inside; dashed line, twofold. (**D**) Fold change of the E-value of the same hit between two calibrations in the benchmark search (per-family medians, hits with E between 0.001 and 10).

Sampling error falls with the square root of the amount of random sequence, so the remedy is to calibrate on more of it. With 16 Mb, ten times the default, the seed-to-seed shift of the threshold fell from 1.0 to 0.35 bits, as sampling theory predicts (Fig. 3B), and E-values of the same hits varied by only 1.08-fold. Against an independent 16-Mb calibration, default calibrations misplaced the threshold by a median of 0.74 bits, the error expected of an unbiased estimate with this scatter, so the longer calibrations are about three times more accurate. For the 260 benchmark models they took 7.2 h on one GPU, about as long as the default calibration takes on 40 CPU cores.

We then recalibrated the whole Rfam library on 16 Mb. For local Inside scores, which cmsearch reports by default, the E-values distributed with Rfam are off by a median of 1.35-fold, and by more than twofold for 12.5% of the 4,227 models; for glocal Inside scores, by more than twofold for 31% (Figs 1A and 3C). The errors are symmetric in sign, as expected of sampling noise.

### 3.4 Sequence filters discard homologs whose similarity lies in base pairing

For search, the GPU first provides an exhaustive reference. On the RMARK benchmark, exhaustive search found 4.2–4.6 percentage points more positives than default Infernal at one false positive per query across three replicates, most of them among test sequences 40–50% identical to the training set (Supplementary Fig. S2). Any filter setting can be replayed on this reference: keeping only the exhaustive hits inside the envelopes that a setting passes reproduced Infernal’s measured sensitivity at that setting to within 0.2 points (Fig. 4B). The filters therefore determine which homologs are found; banding and scoring in the CM stages make little difference. A factorial experiment located the losses. Opening the early filters (SSV, and Viterbi with its composition-bias filter) or loosening the later Forward stages each recovered about a third of the gap to exhaustive search, and only opening both recovered nearly all of it, while sending 17 times more sequence to the CM stages (Fig. 4B).

**Fig. 4.**
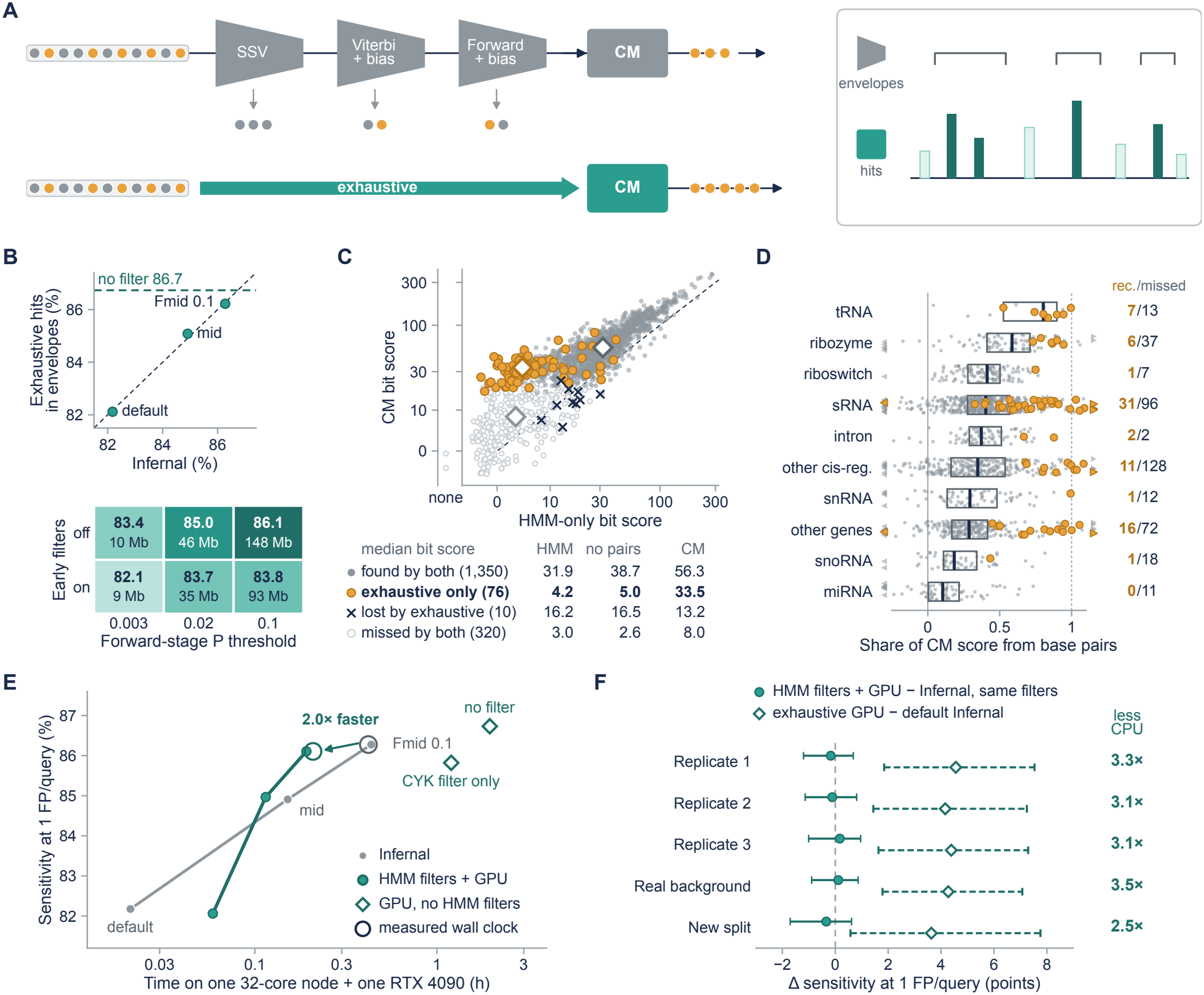
Sequence filters discard homologs whose similarity lies in base pairing; behind permissive filters, the GPU recovers them with much less CPU work. (**A**) Infernal’s profile-HMM filters pass only part of the target to the CM; exhaustive GPU search scores all of it, and a filter setting is replayed by keeping the exhaustive hits inside the envelopes it passes. (**B**) Replayed against measured sensitivity at one false positive per query for three Infernal settings (RMARK, 260 families, 1,756 positives; dashed, exhaustive search), and sensitivity with the early filters on or off and three Forward-stage thresholds, with the sequence passed to the CM stages. (**C**) Bit scores of each positive with its family’s profile HMM and CM, by whether default Infernal and exhaustive search find it at E ≤ 0.01; table, median scores with the HMM, a CM without base pairs and the full CM. Eight of the ten positives found only by default Infernal are longer than the maximum hit length W, which bounds the bands of the filter-free mode. (**D**) Share of each positive’s CM score contributed by base pairs, by RNA class (boxes, median and interquartile range; amber, recovered only by exhaustive search; triangles, beyond the axis); right, recovered over missed by default Infernal. (**E**) Sensitivity against time on one 32-core server with one RTX 4090 for Infernal at three filter settings, the same filters followed by GPU CM computation, and GPU search without HMM filters, with the CYK filter (filter-free) and without it (exhaustive); hollow circles, measured wall clock. (**F**) GPU CM computation behind the most permissive filters minus Infernal with the same filters (filled) and exhaustive GPU search minus default Infernal (open), with paired 95% intervals; right, ratio of CPU work.

To ask why these homologs are lost, we scored every positive without filters with its family’s profile HMM, which sees only sequence, and with its CM (Fig. 4C). The 76 positives that exhaustive search found and default Infernal missed had a median HMM score of 4.2 bits, indistinguishable from chance, but a median CM score of 33.5 bits. A structure-free CM built from the same alignment, identical to the full model except for its base pairs, scored them at a median of 5.0 bits, no better than the HMM, and in 74 of the 76 the base pairs contributed more than all other differences between the models together.

How much a family relies on base pairing varies across RNA classes, and the losses follow this variation (Fig. 4D). Base pairs supplied a median of 10% of the CM score of microRNA homologs and 18% for snoRNAs, and exhaustive search recovered almost none of the homologs of these classes that default Infernal missed. For sRNAs, ribozymes and tRNAs, base pairs supplied 40–80% of the score, and these classes accounted for most of the recovered homologs; in tRNA, exhaustive search recovered seven of the 13 positives that default Infernal missed.

### 3.5 Behind permissive filters, Infernal-GPU recovers lost homologs with a third of the CPU work

Infernal’s most permissive filters already pass nearly all of the homologs that exhaustive search adds (Fig. 4B), but they flood the CM stages with sequence, which then take about 80% of Infernal’s CPU work. We therefore placed the GPU behind the filters: Infernal writes out the envelopes that pass its HMM stages, the GPU scores them, and Infernal rescans the candidates (Section 2.5). Behind the filters of `--mid --Fmid 0.1`, this pipeline reached 86.1% sensitivity at one false positive per query, indistinguishable from Infernal with the same filters (86.3%; paired 95% interval of the difference −1.2 to 0.7 points) and close to exhaustive search (86.7%), while cutting CPU work 3.3-fold, from 13.9 to 4.2 core-hours, for 3.7 GPU-minutes. On one 32-core server with one GPU it finished in half of Infernal’s time (Fig. 4E); the GPU was busy for a small part of the run, and the HMM filters and rescans on the CPU now take most of the time. The result held on two further replicates and two independently constructed benchmarks, with 2.5- to 3.5-fold less CPU work across all five (Fig. 4F); the saving grows as the filters open, because opening them moves work into the CM stages.

## 4 Discussion

Infernal owes its speed to two choices, calibrating on little random sequence and letting sequence filters decide where its covariance models look, both made because CM computation was expensive. With the insert-left self-loop evaluated as a prefix computation, in the max-plus semiring for CYK and the log-sum-exp semiring for Inside, and with storage that follows the bands, one GPU does the CM work of hundreds of CPU cores in calibration and search alike, while Infernal still computes every final score. Both choices can therefore be revisited.

With calibration taking seconds to minutes per model, short calibrations are no longer necessary. The E-values of current models carry a sampling error large enough to move true homologs across the inclusion threshold from one calibration to the next, and for one Rfam model in eight they are off by more than twofold. Such errors matter most where E-values gate decisions automatically, as in iterative searches and in the alignments built for each query in RNA structure prediction, which are also the settings in which a model must be calibrated on the spot. We provide Rfam 15.0 recalibrated on ten-fold longer random sequence and suggest that model libraries be calibrated this way at release.

Sequence filters trade sensitivity for speed, and an unfiltered reference makes it possible to measure both how much sensitivity is lost and why. The homologs lost are those for which CMs were designed: sequences that have drifted to the level of chance while keeping their base pairs, concentrated in structured classes such as tRNAs, ribozymes and many bacterial sRNAs. Recovering them requires opening the filters and accepting much more CM computation, which the GPU absorbs; moving the HMM filters and final rescans to the GPU as well would remove the CPU stages that now dominate. GPU sequence prefilters such as Riboseek (Park *et al*. 2026) accelerate the other end of the same pipeline, so the two approaches are complementary. Beyond Infernal, the prefix formulation applies to the insert states of other profile grammars and to the Inside–Outside computations of CM alignment.

## Data availability

Infernal-GPU is available at https://github.com/void-echo/infernal-gpu and from PyPI; Rfam 15.0 models recalibrated on 16 Mb of random sequence are attached to its v0.1.0 release. Rfam 15.0 models and seed alignments are available from the Rfam database; genome sequences from NCBI (NC_013790.1, NC_000913.3) and the human gut metagenome assembly GCA_009685265.1. The benchmark constructions, calibration tables, search out-puts, timing logs and the scripts that reproduce every figure will be deposited at Zenodo.

## References

Blelloch GE. Prefix Sums and Their Applications. Technical Report CMU-CS-90-190. School of Computer Science, Carnegie Mellon University, 1990.

Durbin R, Eddy SR, Krogh A et al. Biological Sequence Analysis: Probabilistic Models of Proteins and Nucleic Acids. Cambridge, UK: Cambridge University Press, 1998.

Eddy SR. Accelerated profile HMM searches. PLoS Comput Biol 2011;7:e1002195.

Eddy SR, Durbin R. RNA sequence analysis using covariance models. Nucleic Acids Res 1994;22:2079–88.

Efron B, Tibshirani RJ. An Introduction to the Bootstrap. New York: Chapman & Hall, 1993.

Freyhult EK, Bollback JP, Gardner PP. Exploring genomic dark matter: a critical assessment of the performance of homology search methods on noncoding RNA. Genome Res 2007;17:117–25.

Ganesan N, Chamberlain RD, Buhler J et al. Accelerating HMMER on GPUs by implementing hybrid data and task parallelism. In: Proceedings of the First ACM International Conference on Bioinformatics and Computational Biology. New York, NY, USA: ACM, 2010, 418–21.

Hassan S, Särkkä S, García-Fernández ÁF. Temporal parallelization of inference in hidden Markov models. IEEE Trans Signal Process 2021;69:4875–87.

Jiang H, Ganesan N. CUDAMPF: a multi-tiered parallel framework for accelerating protein sequence search in HMMER on CUDA-enabled GPU. BMC Bioinformatics 2016;17:106.

Johnson M. Parsing in parallel on multiple cores and GPUs. In: Molla D, Martinez D (eds), Proceedings of the Australasian Language Technology Association Workshop 2011, Canberra, Australia. Australasian Language Technology Association, 2011, 29–37.

Kallenborn F, Chacon A, Hundt C et al. GPU-accelerated homology search with MMseqs2. Nat Methods 2025;22:2024–7.

Kalvari I, Nawrocki EP, Ontiveros-Palacios N et al. Rfam 14: expanded coverage of metagenomic, viral and microRNA families. Nucleic Acids Res 2021;49:D192–200.

Khajeh-Saeed A, Poole S, Perot JB. Acceleration of the Smith–Waterman algorithm using single and multiple graphics processors. J Comput Phys 2010;229:4247–58.

Klein RJ, Eddy SR. RSEARCH: finding homologs of single structured RNA sequences. BMC Bioinformatics 2003;4:44.

Kogge PM, Stone HS. A parallel algorithm for the efficient solution of a general class of recurrence equations. IEEE Trans Comput 1973;C-22:786–93.

Kolbe DL, Eddy SR. Fast filtering for RNA homology search. Bioinformatics 2011;27:3102–9.

Ladner RE, Fischer MJ. Parallel prefix computation. J ACM 1980;27:831–8.

Leahy SC, Kelly WJ, Altermann E et al. The genome sequence of the rumen methanogen Methanobrevibacter ruminantium reveals new possibilities for controlling ruminant methane emissions. PLoS One 2010;5:e8926.

Lei G, Dou Y, Wan W et al. CPU-GPU hybrid accelerating the Zuker algorithm for RNA secondary structure prediction applications. BMC Genomics 2012;13:S14.

McVicar N, Ruzzo WL, Hauck S. Accelerating ncRNA homology search with FPGAs. In: Proceedings of the ACM/SIGDA International Symposium on Field Programmable Gate Arrays. New York, NY, USA: ACM, 2013, 43–52.

Miladi M, Sokhoyan E, Houwaart T et al. GraphClust2: annotation and discovery of structured RNAs with scalable and accessible integrative clustering. GigaScience 2019;8:giz150.

Moscola J, Cytron RK, Cho YH. Hardware-accelerated RNA secondary-structure alignment. ACM Trans Reconfigurable Technol Syst 2010;3:14.

Nawrocki EP, Eddy SR. Query-dependent banding (QDB) for faster RNA similarity searches. PLoS Comput Biol 2007;3:e56.

Nawrocki EP, Eddy SR. Infernal 1.1: 100-fold faster RNA homology searches. Bioinformatics 2013;29:2933–5.

Nawrocki EP, Eddy SR. Infernal User’s Guide, Version 1.1.5. 2023. http://eddylab.org/infernal/Userguide.pdf (9 October 2026, date last accessed).

Nawrocki EP, Kolbe DL, Eddy SR. Infernal 1.0: inference of RNA alignments. Bioinformatics 2009;25:1335–7.

Ontiveros-Palacios N, Cooke E, Nawrocki EP et al. Rfam 15: RNA families database in 2025. Nucleic Acids Res 2025;53:D258–67.

Park S, Didi K, Favor A et al. Fast remote nucleotide sequence alignment with Riboseek. bioRxiv, 10.64898/2026.07.31.741718, 2026, preprint: not peer reviewed.

Rizk G, Lavenier D. GPU accelerated RNA folding algorithm. In: Allen G, Nabrzyski J, Seidel E et al. (eds), Computational Science – ICCS 2009. Lecture Notes in Computer Science, Vol. 5544. Berlin, Heidelberg: Springer, 2009, 1004–13.

Sakakibara Y, Brown M, Hughey R et al. Stochastic context-free grammars for tRNA modeling. Nucleic Acids Res 1994;22:5112–20.

Schmidt B, Kallenborn F, Chacon A et al. CUDASW++4.0: ultra-fast GPU-based Smith–Waterman protein sequence database search. BMC Bioinformatics 2024;25:342.

Shen T, Hu Z, Sun S et al. Accurate RNA 3D structure prediction using a language model-based deep learning approach. Nat Methods 2024;21:2287–98.

Stern HA, Mathews DH. Accelerating calculations of RNA secondary structure partition functions using GPUs. Algorithms Mol Biol 2013;8:29.

Wang W, Feng C, Han R et al. trRosettaRNA: automated prediction of RNA 3D structure with trans-former network. Nat Commun 2023;14:7266.

Weinberg Z, Ruzzo WL. Sequence-based heuristics for faster annotation of non-coding RNA families. Bioinformatics 2006;22:35–9.

Weinberg Z, Lünse CE, Corbino KA et al. Detection of 224 candidate structured RNAs by comparative analysis of specific subsets of intergenic regions. Nucleic Acids Res 2017;45:10811–23.

Xia F, Dou Y, Zhou D et al. Fine-grained parallel RNA secondary structure prediction using SCFGs on FPGA. Parallel Comput 2010;36:516–30.

Yao Z, Weinberg Z, Ruzzo WL. CMfinder—a covariance model based RNA motif finding algorithm. Bioinformatics 2006;22:445–52.

Yi Y, Lai C-Y, Petrov S. Efficient parallel CKY parsing using GPUs. J Log Comput 2014;24:375–93.

Zhang T, Singh J, Litfin T et al. RNAcmap: a fully automatic pipeline for predicting contact maps of RNAs by evolutionary coupling analysis. Bioinformatics 2021;37:3494–500.

Zhang C, Zhang Y, Pyle AM. rMSA: a sequence search and alignment algorithm to improve RNA structure modeling. J Mol Biol 2023;435:167904.

