## Supplementary Material for "Infernal-GPU: accurate E-values and sensitive covariance-model search for structured RNAs"

### Supplementary Tables

**Table S1. Agreement of Infernal-GPU with Infernal.** Calibrations are compared by the bit score that gives E = 0.01 in a 100-Mb search; search hits count as identical when coordinates, bit score and E-value agree. Filter-free search reproduces `cmsearch --nohmm` and exhaustive search `cmsearch --nohmm --noF6`, both in non-truncated mode; the RMARK benchmark is replicate 1, reported to E ≤ 10.

| Comparison | Result |
|---|---|
| Calibration, 260 benchmark models, GPU versus cmcalibrate on identical random sequence | median threshold difference 0.023 bits (four modes); between random seeds, 1.0 bit |
| Calibration, 4,227 Rfam 15.0 models, GPU versus distributed parameters | median difference 0.023 bits; 95% of model–mode pairs within 0.1 bits |
| Calibration, 260 benchmark models, RTX 4090 versus A100 | 1,040 of 1,040 model–mode parameter sets bit-identical |
| Search, 4,227 Rfam models × *M. ruminantium* genome, filter-free | 129 of 129 hits identical (91 of 91 with E < 0.01) |
| Search, 38 Rfam models stratified by cost, including the two largest rRNA models, same genome, filter-free | final hits identical for 38 of 38 models |
| Search, eight models × *E. coli* K-12 genome, filter-free | 105 of 105 hits identical (100 of 100 with E < 0.01) |
| Search, RMARK benchmark, 260 families, filter-free | 4,433 of 4,433 hits identical |
| Search, RMARK benchmark, 260 families, exhaustive | 4,948 of 4,948 hits identical |

**Table S2. Calibration wall clock on one server.** cmcalibrate with `--cpu 32` on a dual-socket AMD EPYC 7742 server against Infernal-GPU (four modes, one RTX 4090). The first seven models are from the RMARK benchmark, the SSU rRNA models from Rfam 15.0.

| Model | cmcalibrate wall (s) | cmcalibrate CPU work (core-s) | Infernal-GPU (s) | Ratio of wall times |
|---|---:|---:|---:|---:|
| SECIS_1 | 24 | 718 | 3.1 | 7.7 |
| tRNA | 50 | 1,514 | 5.6 | 8.9 |
| U1 | 82 | 2,512 | 8.8 | 9.3 |
| 6S | 172 | 5,357 | 14.7 | 11.8 |
| Cobalamin | 319 | 9,998 | 25.4 | 12.6 |
| tmRNA | 372 | 11,466 | 35.9 | 10.4 |
| RNaseP_bact_a | 809 | 24,629 | 63.2 | 12.8 |
| SSU_rRNA_archaea | 2,055 | 59,627 | 253.7 | 8.1 |
| SSU_rRNA_bacteria | 2,244 | 65,451 | 280.5 | 8.0 |

### Supplementary Figures

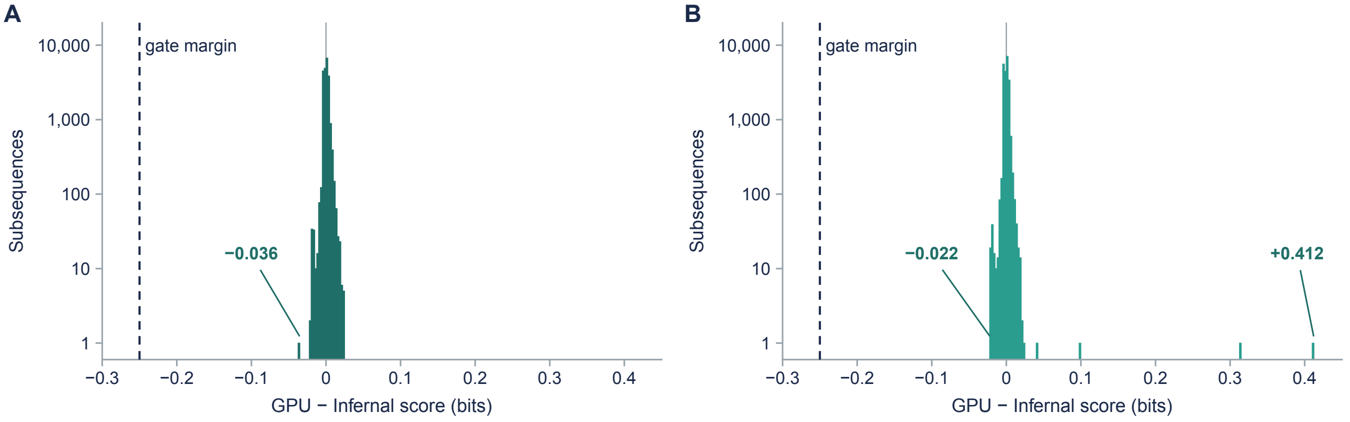


**Fig. S1. GPU scores near the gating thresholds.** Distribution of GPU minus Infernal CYK (**A**) and Inside (**B**) scores for 21,861 subsequences near the gating thresholds of an eight-model search of the *M. ruminantium* genome, computed with Infernal's own scoring routines. The largest GPU underestimates, 0.036 bits for CYK and 0.022 bits for Inside, are an order of magnitude smaller than the 0.25-bit margin by which the gates are lowered (dashed). Inside differences are larger than CYK differences because Infernal evaluates log-sums with a lookup table, whereas the GPU computes them in floating point; the largest positive differences occur on low-complexity, AU-rich subsequences and can only admit extra candidates, which Infernal's rescans score exactly.

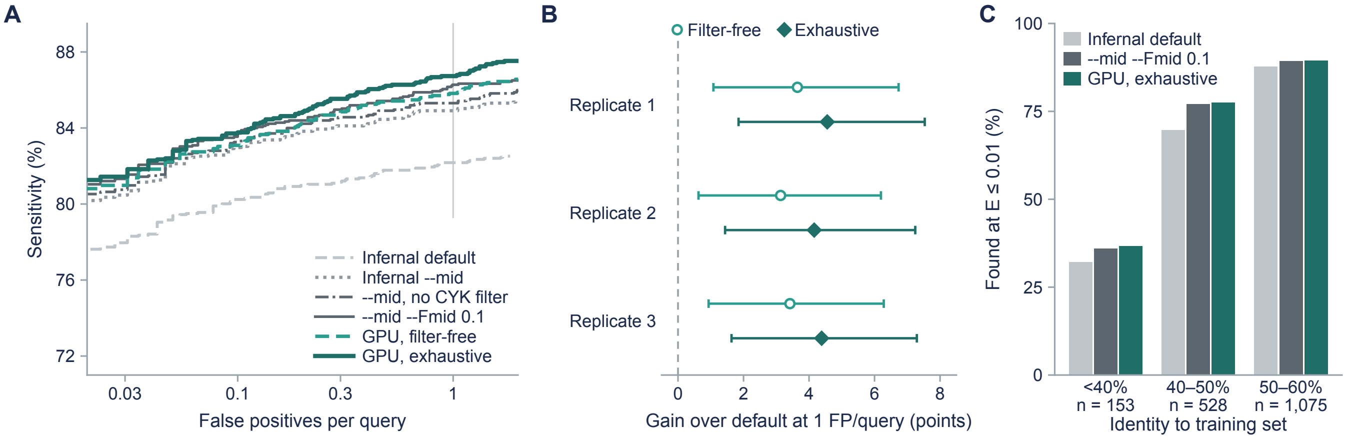


**Fig. S2. Remote-homolog recovery on the RMARK benchmark.** (**A**) Sensitivity against false positives per query (260 families, 1,756 positives) for Infernal at default, `--mid`, `--mid` without its CYK filter (`--noF6`) and `--mid --Fmid 0.1`, and for GPU search without HMM filters, with the CYK filter (filter-free) and without it (exhaustive). (**B**) Gain of filter-free and exhaustive GPU search over default Infernal at one false positive per query on three benchmark replicates, with paired family-bootstrap 95% intervals. (**C**) Positives found at $E \leq 0.01$ by default Infernal, Infernal with `--mid --Fmid 0.1` and exhaustive GPU search, grouped by their highest identity to the training sequences. The vertical line in **A** marks one false positive per query.
